# IFI207 promotes antiviral responses by modulating STING ubiquitination and degradation

**DOI:** 10.64898/2026.03.05.709838

**Authors:** Takuji Enya, Wenming Zhao, Geetanjali, Bin He, Susan R Ross

## Abstract

Aim2-like receptors (ALRs) play crucial roles in innate immune signaling pathways and demonstrate strong positive selection likely driven by pathogens. IFI207, an ALR found in all *Mus* species, enhances interaction with and stabilization of STING, contributing to the control of Murine Leukemia Virus (MLV) infection. We show here that IFI207 enhances the type 1 interferon response by inhibiting activation-induced K63-linked ubiquitination of STING, thereby preventing its recognition by hepatocyte growth factor-regulated tyrosine kinase substrate (HRS), a key component of the ESCRT complex, and its subsequent degradation in lysosomes. IFI207 promotes downstream signaling in the STING pathway in multiple cell types and moreover enhances the STING-dependent response to herpesvirus simplex 1 infection ex vivo and in vivo. We also show that IFI207 likely functions in dendritic cells to suppress MLV infection. Our study reveals that IFI207 acts as a modulator in the STING pathway, strengthening the host’s defense against viral infections and suggests that the expansion of the *Alr* locus in mice may have occurred in response to endemic viruses.

## Introduction

The innate immune system detects pathogen-associated molecular patterns (PAMPs) through pattern recognition receptors (PRRs) such as Toll-like receptors (TLRs), retinoic acid-inducible gene I-like receptors (RLRs), Absent in Melanoma 2 (AIM2)-like receptors (ALRs), cyclic GMP-AMP (cGAMP) synthase (cGAS) and DEAD box helicases (DDXs). Upon ligand binding, these PRRs trigger transcription and production of type I interferons (IFN-I) and other cytokines (1–5). IFN-I in turn activates the JAK-STAT pathway via the IFN-alpha/beta receptor (IFNAR), inducing expression of IFN-stimulated genes (ISG) that play critical roles in antiviral defense. The stimulator of interferon genes (STING) protein is downstream of several PRRs that bind nucleic acids, including cGAS, ALRs and DDXs, and thus plays a crucial role in protecting the host against pathogens (6–8). For example, cGAS recognizes DNA generated during virus infection and synthesizes cGAMP, which binds to STING on the endoplasmic reticulum (ER), triggering its activation and subsequent dimerization (9). The STING dimers associate with TANK-binding kinase 1 (TBK1), become phosphorylated and translocate to the Golgi apparatus. TBK1 also phosphorylates transcription factors, including interferon regulatory factor (IRF) 3, which moves to the nucleus, leading to the production of IFN-I and pro-inflammatory cytokines (6). After activation, STING is ubiquitinated and degraded, either in the lysosome or in autophagosomes (10, 11).

STING activation is regulated by various types of modifications, including phosphorylation and ubiquitination (12–14). Ubiquitin regulates cellular processes through proteasome-dependent and -independent mechanisms and is important for STING-mediated antiviral signaling (14). There are seven types of lysine (K) linkages on STING that can be ubiquitinated, K6, K11, K27, K29, K33, K48, and K63 (15). Among these, K48 and K63 linkages are believed to be the main types of ubiquitination that regulate STING’s activity. TRIM32 and TRIM56 mediate K63-linked polyubiquitination of STING, facilitating TBK1 recruitment, thereby leading to antiviral responses (16, 17). K63-ubiqutination of amino acid 288 also occurs during STING activation and is followed by its degradation (11). K48-linked ubiquitination by TRIM29 and TRIM30α also leads to STING degradation (18). Other host proteins such as the deubiquitinase CYLD interact with and selectively remove K48-linked polyubiquitin chains on STING, by this means boosting the innate antiviral response (19).

Proteins are transported for degradation by the endosomal complexes required for transport (ESCRT) machinery, which consists of five distinct complexes (ESCRT-0, -I, -II, -III, and Vps4), each with specific roles in recognizing ubiquitinated membrane proteins, membrane deformation, and abscission (20). Ubiquitinated proteins are recognized by the ESCRT-0 complex protein hepatocyte growth factor-regulated tyrosine kinase substrate (HRS) (19). HRS recognizes ubiquitinated STING and promotes its degradation, thereby terminating STING-mediated signaling (10, 21).

ALRs are a family of proteins with evidence of pathogen-driven strong positive selection that have important roles in innate immune signaling pathways (22–25). ALRs have an N-terminal pyrin domain (PYD) that facilitates homotypic and heterotypic interactions with PYD-containing and other proteins, and a C-terminal hematopoietic interferon-inducible nuclear (HIN) domain, which binds DNA (4, 24, 26). *ALR*s are found in mammalian species at a single genetic locus, and the locus has dramatically expanded in mice (22–25). We reported that IFI207 is unique among ALRs due to a large repeat region consisting of 14 amino acid S/T-rich repeats separating the N-terminal PYD and C-terminal HIN domains that is predicted to form a coiled domain (23). Unlike other ALRs that bind nucleic acid, IFI207 binds to and stabilizes STING protein, and the repeat region plays a crucial role in facilitating this interaction. This stabilization contributes to heightened innate responses to STING agonists, as well as the control of MLV infection *in vivo*. Moreover, basal STING levels are reduced in the absence of IFI207.

Here we asked how IFI207 contributes to STING stabilization. We show that that IFI207 reduces STING degradation via acidic compartments by blocking its K63 ubiquitination. Using bone marrow-derived macrophages (BMDMs) and fibroblasts isolated from IFI207 knockout (KO) and wild-type (WT) mice, we show that by diminishing K63 ubiquitination and degradation, IFI207 causes increased STING activation and increased type I interferon induction in response to both the STING agonist DMXAA and HSV-1. Because IFI207 reduced K63-linked ubiquitination on STING, binding of HRS to STING following stimulation with the STING agonist DMXAA or herpes simplex virus (HSV)-1 was reduced in IFI207-positive cells. As a result, IFI207 KO mice are more susceptible to both HSV-1 and MLV. These findings suggest that IFI207 blocks STING’s degradation via the ESCRT pathway following its activation and that this enhances its antiviral activity.

## Results

### IFI207 reduces STING degradation via ubiquitination

STING induction, trafficking and degradation require ubiquitination, V-ATPase function and acidified endolysosomes (10, 27). TAK243, a ubiquitin activating enzyme inhibitor, enhances STING signaling by inhibiting STING ubiquitination and degradation, resulting in increased phosphorylated STING (pSTING), pTBK1 and pIRF3 (10). Bafilomycin A1 (BafA1), which inhibits the endosomal V-ATPase and thus acidification of late endosomes and lysosomes, also reduces STING degradation, suggesting that lysosomes are important for trafficking-dependent STING degradation (27).

We first examined whether IFI207 inhibited STING degradation via the lysosome and was dependent on ubiquitination, using BMDMs and fibroblasts isolated from IFI207 KO and WT mice. STING RNA levels were equivalent in IFI207 and WT KO cells, as we previously reported (Suppl. Fig. 1A and 1B) (23). However, total basal STING protein levels were reduced in IFI207 KO BMDMs and fibroblasts over 20% and 30%, respectively (Fig. 1A and 1C). Cells were pre-treated with either TAK243 or BafA1, stimulated with DMXAA, and STING and p-STING levels were quantified by western blots. By 1 hour (hr) after stimulation, total STING levels were reduced compared to untreated cells for both WT and IFI207 KO cells, as were DMXAA-induced p-STING levels in IFI207 KO cells. In macrophages, TAK243 blocked the decrease in total and p-STING levels that occurred at 1 and 3 hrs post-DMXAA treatment in both WT and 207 KO cells (Fig. 1A). Indeed, p-STING were higher in TAK243-treated WT and IFI207 KO macrophages than they were with DMXAA alone (Suppl. Fig. 1C). IFNβ RNA, which increased in both WT and KO cells at 2 hr post-induction, was partially restored by treatment with TAK23 but not bafA, particularly in WT cells as has been previously shown (Fig. 1B) (10). In BafA1-treated cells, the levels of total and p-STING were stabilized in the KO cells, although p-STING levels were reduced compared to cells treated with DMXAA alone (Fig. 1A). Notably, while TAK243 significantly induced IFNβ RNA levels over 3-fold in WT and IFI207 KO macrophages after DMXAA treatment, BafA1 did not (Fig. 1B and Suppl. 1C).

**Fig. 1.**
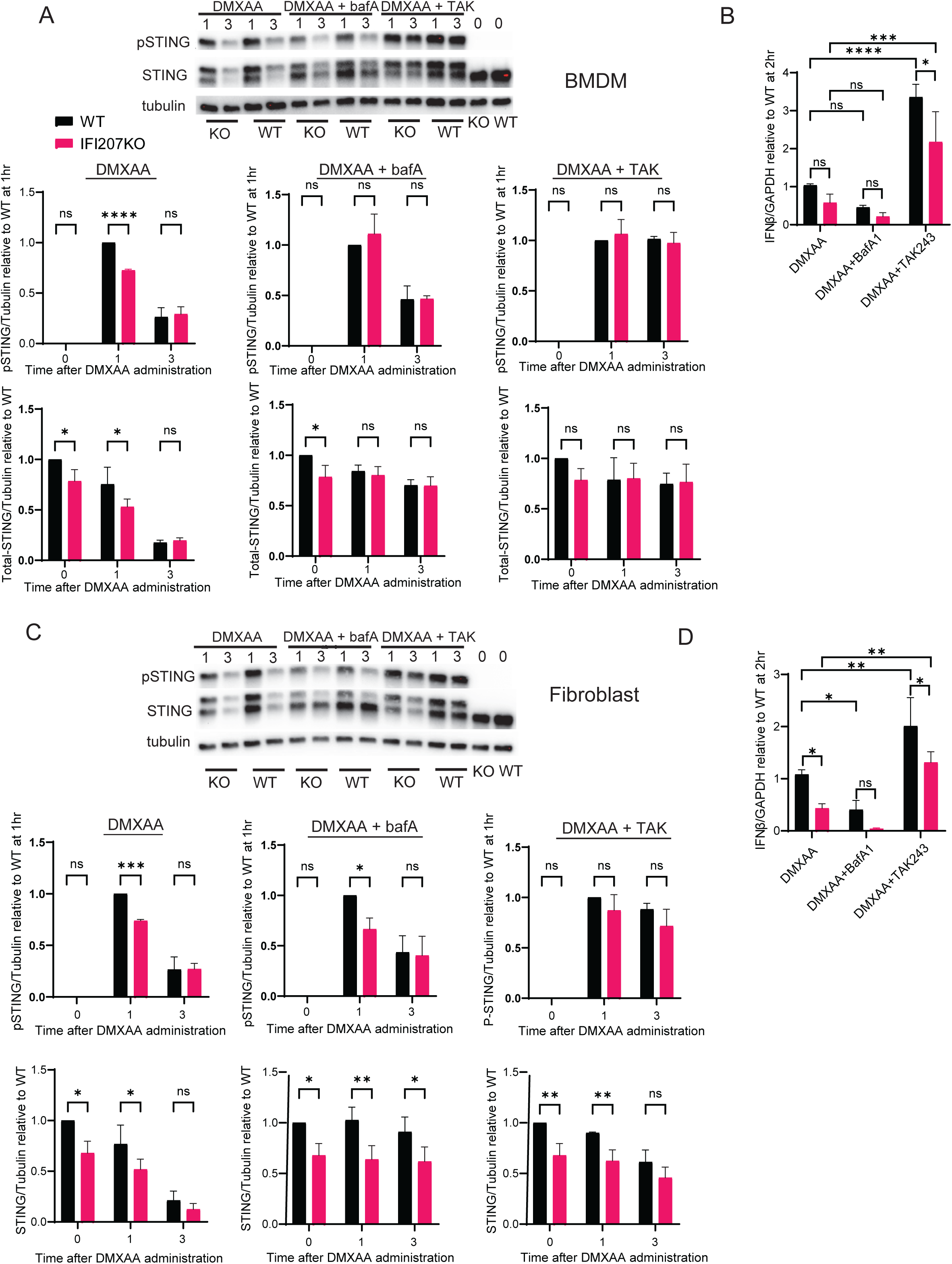
IFI207 reduces STING degradation via blocking ubiquitination. BMDMs (A, B) and fibroblasts (C, D) from the indicated mice were pre-treated with either bafA1 or TAK243, a ubiquitination inhibitor, and then stimulated with DMXAA for 1 and 3 hr. A) and C) STING and pSTING protein levels were examined by western blot. Shown below is quantification of the average of 3 independent experiments ± SD. B) and D) RT-qPCR was used to determine IFNβ levels at 2 hr post-DMXAA. Means from 3 independent experiments performed with duplicate or triplicate samples per treatment were plotted. Two-way ANOVA was used to determine significance. ****, *P*≤0.0001;***, *P*≤0.0002; **, *P*≤0.007; *, *P*≤0.01; ns, not significant.

Similar to macrophages, TAK243 treatment of DMXAA-stimulated fibroblasts increased the levels of p-STING and stabilized total STING levels over time compared to DMXAA alone (Fig. 1C). However, the difference in total STING and p-STING levels between WT and IFI207 KO cells was maintained. BafA1 treatment also stabilized STING but not p-STING levels in fibroblasts at 3 hr post-DMXAA treatment (Fig. 1C). TAK243 but not BafA1 induced IFNβ RNA (by 1.9-fold in WT and 3-fold in KO cells) and protein (by 1.6-fold in WT and 1.3-fold in KO cells) levels compared to DMXAA treatment alone (Fig. 1D; Suppl. 1D), whereas IFNβ RNA and protein levels decreased in DMXAA/BafA1-treated macrophages and fibroblasts compared to DMXAA-alone treatment (Figs. 1B and 1D). Collectively, these data suggested that IFI207 decreases STING degradation after ubiquitination, thereby enhancing the type I IFN response.

### IFI207 interacts with HRS in the presence of STING

After activation, and ubiquitination, STING is targeted to the lysosome where it is degraded, leading to termination of signaling (10, 21). Proteomic analysis coupled with genetic screening identified HRS and VPS37A, components of the ESCRT machinery, as essential proteins that interact with STING to regulate its trafficking (21).

We next tested whether IFI207 stabilizes STING by preventing its interaction with HRS. As we do not have anti-IFI207 antisera, we transfected HEK293T cells, which are STING-negative, with FLAG-tagged STING and V5-tagged IFI207 plasmids to study their interaction. Twenty-four hrs post-transfection, endogenous HRS was immunoprecipitated from cell extracts, followed by western blotting with the indicated antibodies. As previously shown, STING and HRS bind to each other (Fig. 2A) (10, 21). Phospho-STING also co-immunoprecipitated with HRS, although it was only a small fraction of the total STING present in transfected cells (Suppl. Fig. 1E). IFI207 also co-immunoprecipitated with HRS (Fig. 2A), but not in the absence of STING (Fig. 2B). This indicates that IFI207 may not directly bind HRS but is co-immunoprecipitated in complex with STING. However, IFI207 did not disrupt the interaction with HRS, although it resulted in higher levels of STING in both unstimulated and DMXAA-stimulated cells (Fig. 2A).

**Fig. 2.**
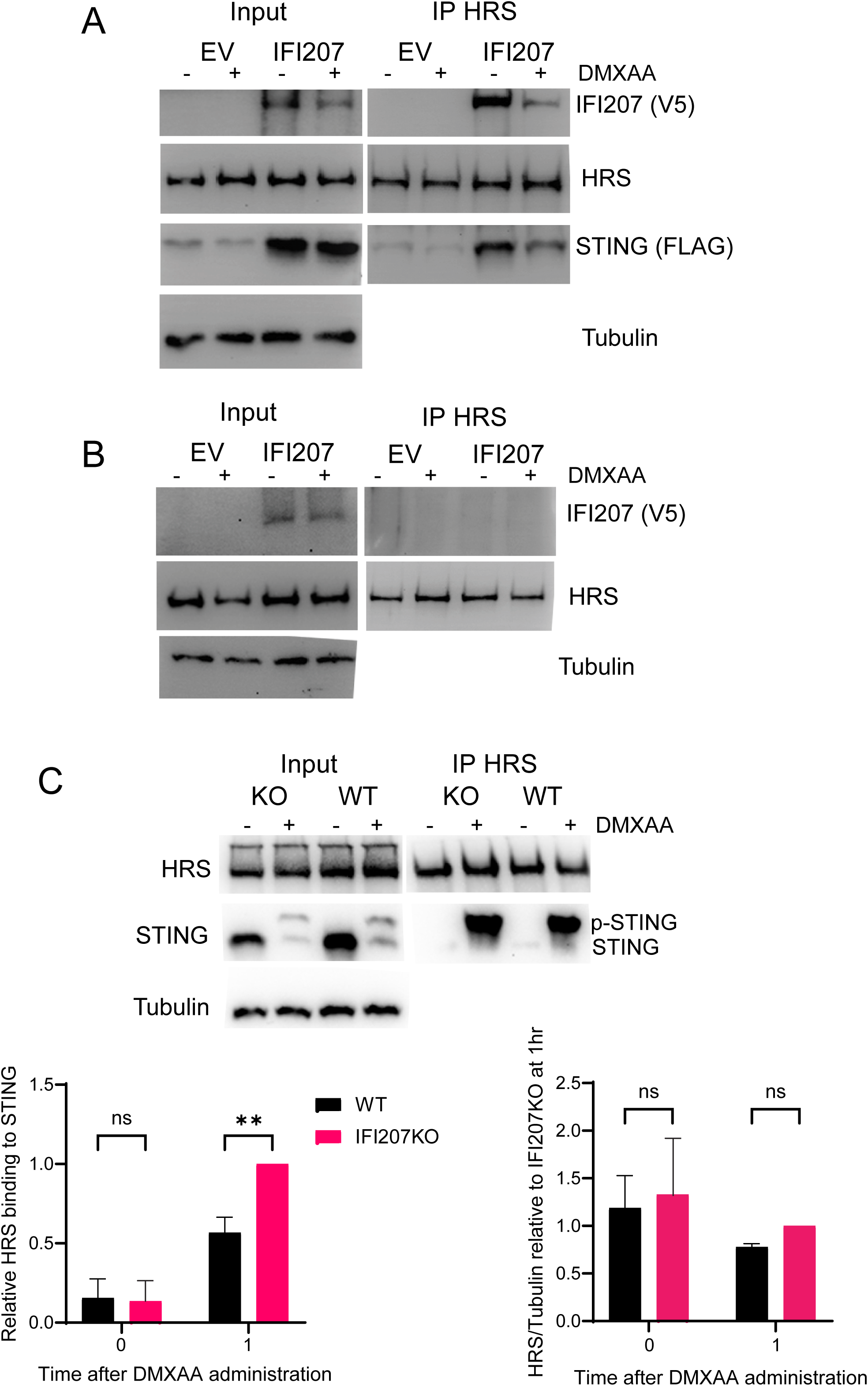
IFI207 interacts with HRS in the presence of STING. A) and B) HEK293T cells were transfected with V5-tagged IFI207 and FLAG-tagged STING expression plasmids (A) or V5-tagged IFI207 alone (B). Twenty four hr after transfection, the cells were treated with DMXAA for 4 h, and then immunoprecipitation and immunoblot analysis were performed with the indicated antibodies. C) BMDMs from the indicated mice were treated with 100 µg/mL DMXAA for 1h. Anti-HRS antibody was used to immunoprecipitate HRS in the cell extracts, and western blots with the indicated anti-HRS and -STING, as well as anti-tubulin antibodies, were performed. Shown below the blots is the average of 3 independent experiments ± SD. Two-way ANOVA was used to determine significance. **, *P*≤0.002; ns, not significant.

We also used BMDMs isolated from WT and IFI207 KO mice to examine the interaction between endogenous STING and HRS. Without stimulation, STING appeared as a single band. In unactivated cells, only very low levels of STING were found in complex with HRS (Fig. 2C). Upon DMXAA stimulation, p-STING co-immunoprecipitated with HRS. In the absence of IFI207, HRS bound greater amounts of STING, although the overall levels of STING were reduced (Fig. 2C). HRS levels were unaffected by DMXAA treatment. These data suggest that endogenous levels of IFI207 prevent the interaction between STING and HRS after activation, which likely contributes to STING stabilization.

We previously showed that the interaction between IFI207 and STING was stabilized by a repeat region unique to IFI207 that is predicted to form a coiled structure (23). Since activated STING is a homodimer, we used AlphaFold analysis to examine the predicted interaction between STING dimers and IFI207 (28). Alphafold predicted that IFI207 interacted with STING in two regions, with the unique coiled structure binding to the N-terminal proximal region (cyan-colored in Supplementary Fig. 2A) and the HINB domain (orange-colored) to the C-terminal region of STING. Modeling of human IFI16, which has a single copy of the S/T-rich repeat and also binds STING, predicted that the pyrin domain (colored green) binds to the C terminal regions of the dimer, but not to the N terminal region (Supplementary Fig. 2B).

### IFI207 reduces K63-linked ubiquitination on STING following DMXAA stimulation

Proteins with K48-linked polyubiquitin chains are degraded by the ubiquitin-proteasome system, whereas K63-linked polyubiquitylation acts as a sorting signal within the endosomal membrane, directing proteins towards lysosomal degradation (29). HRS specifically binds proteins with K63 ubiquitin linkages and recruits them via the ESCRT complex to acidic compartments (30, 31).

We next investigated whether IFI207 altered K63 linked ubiquitination of STING. BMDMs isolated from IFI207 KO and WT were treated with DMXAA for 1 hr and an anti-K63-linkage-specific polyubiquitin antibody was used to immunoprecipitate the cell extracts. The immunoprecipitates were analyzed by western blotting using anti-STING antibodies. In the absence of DMXAA, very low levels of STING were precipitated with the anti-K63 Ub antibodies from either WT or KO cells (Fig. 3A). Upon stimulation, anti-K63 Ub strongly immunoprecipitated p-STING, and although STING levels were lower in the KO cells, more ubiquitinated pSTING was precipitated than from WT cells (Fig. 3A). IFI207 thus protects activated STING from K63-linked ubiquitination.

**Fig. 3.**
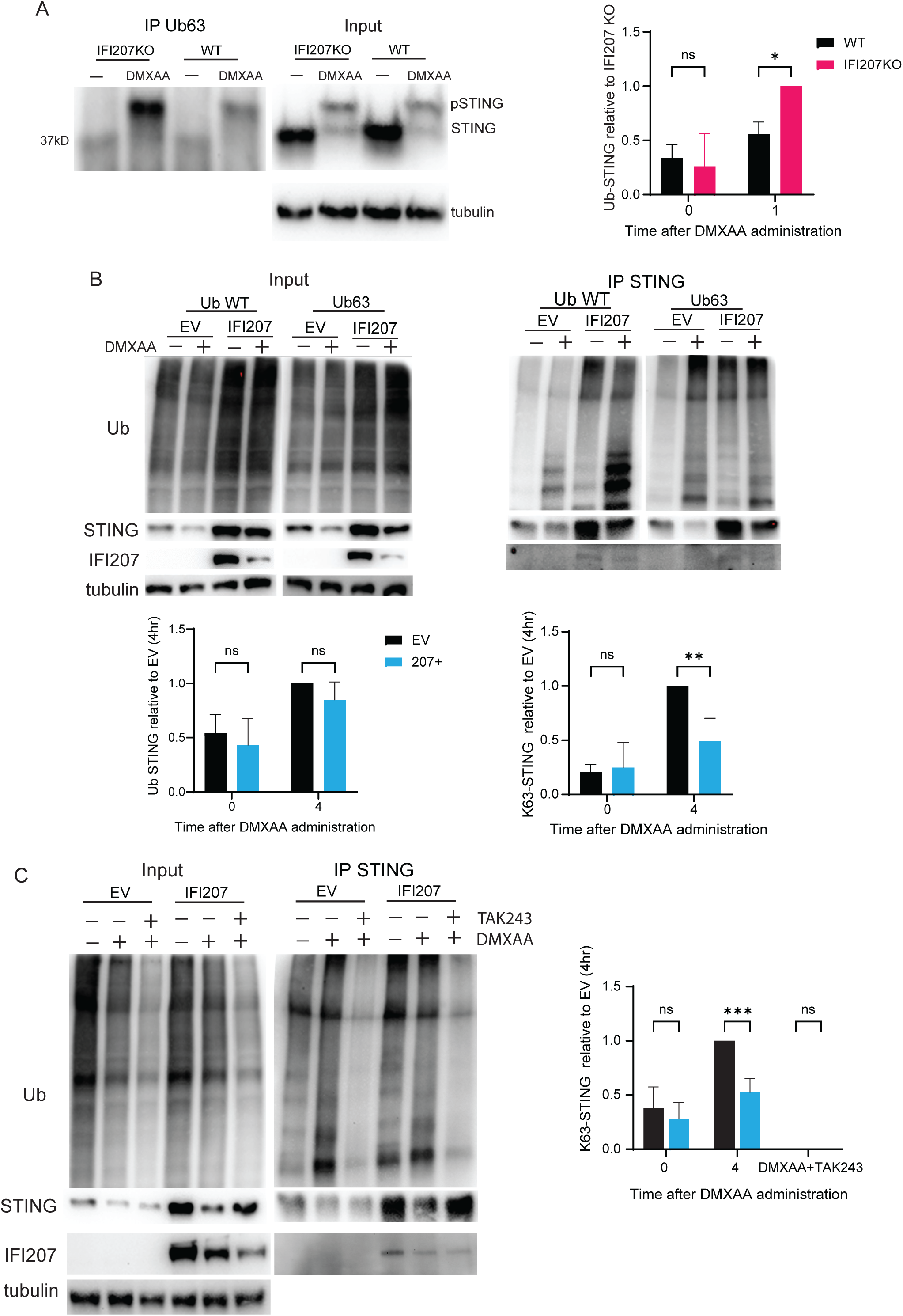
IFI207 reduces K63-linked ubiquitination on STING following DMXAA stimulation. A) BMDMs from the indicated mice were treated with 100 µg/mL DMXAA for 1 hr. An anti-K63-linkage-specific polyubiquitin antibody was used to immunoprecipitate cell extracts. The immunoprecipitates were then analyzed by western blotting using anti-STING and anti-tubulin antibodies. Shown to the right is quantification of 3 independent experiments ± SD. Two-way ANOVA was used to determine significance. **, *P*≤0.03; ns, not significant. B) HEK293T cells were transfected with either HA-tagged Ub-WT or -K63 expression plasmids along with Flag-tagged STING and V5-tagged IFI207. 24 hr after transfection, the cells were treated with 100 μg/ml DMXAA for 2 hr. Immunoprecipitation and immunoblot analysis were performed with the indicated antibodies. To quantify, the ubiquitin signal was normalized to the STING signal and then the EV (4 hr) sample was set to 1. Shown below is quantification of the average of 3 independent experiments ± SD. Two-way ANOVA was used to determine significance. *, *P*≤0.01; ns, not significant. C) HEK293T cells were transfected with HA-tagged Ub-63 plasmids along with Flag-tagged STING and V5-tagged IFI207. 24 hr after transfection, the cells were pre-treated with TAK243 for 30 mins and then stimulated with DMXAA for 4 hr. Immunoprecipitation and immunoblot analysis were performed with the indicated antibodies. Shown to the right is quantification of the average of 3 independent experiments ± SD. Two-way ANOVA was used to determine significance. **, *P*≤0.0008; ns, not significant.

To further confirm that IFI207 reduces K63-linked ubiquitination of STING, we co-transfected IFI207 and STING expression plasmids with either HA-Ub (wild-type) or HA-Ub (K63-specific) expression plasmids into HEK293T cells and examined the effect of IFI207 on STING ubiquitination. Although there was increased ubiquitination of STING using HA-Ub (wild-type), no significant differences were observed in the presence or absence of IFI207 (compare empty vector (EV) to IFI207 lanes in Fig. 3B). However, the levels of K63-linked ubiquitination on STING were significantly reduced in HA-Ub (K63)-transfected cells in the presence of IFI207 (Fig. 3B), further indicating that IFI207 blocks K63-linked ubiquitination of STING.

TAK243 treatment, which blocks the conjugation of ubiquitin to target sites, stabilized STING levels after DMXAA activation (Fig. 1). To show that this was the direct result of inhibiting K63-linked STING ubiquitination, we also treated IFI207/STING-transfected cells with TAK243. TAK243 inhibited global ubiquitination in DMXAA-treated HEK293T cells and degradation of STING (Fig. 3C). TAK243 also reduced K63-ubiquitination of STING after DMXAA treatment (Fig. 3C). Interestingly, IFI207 levels were diminished upon DMXAA treatment, but TAK243 did not restore its levels to that seen in untreated cells, indicating that IFI207 is degraded via a non-ubiqutin-dependent pathway (Fig. 3C). Thus, IFI207 reduces K63-linked ubiquitination on STING, thereby preventing STING’s degradation via the ESCRT pathway.

### IFI207 enhances the type I IFN response to HSV-1

STING protein levels and signaling are decreased in IFI207 KO bone-marrow derived macrophages (BMDMs) and fibroblasts (Fig. 1) (23). Similar results were seen in IFI207 KO bone-marrow derived dendritic cells (BMDCs) (Supplementary Fig. 3). During its activation, STING is phosphorylated by TBK1, which then trans-phosphorylates other TBK1 molecules and IRF3 and NF-κB. These transcription factors move to the nucleus, leading to the production of IFN-I and pro-inflammatory cytokines (6). If STING degradation was suppressed by its association with IFI207, P-TBK1 and P-IRF3 levels should be enhanced by IFI207. To confirm this, BMDMs (Fig. 4A) and BMDCs (Fig. 4B) were isolated from IFI207KO and WT mice and then treated with DMXAA. IFI207 increased p-TBK1 and p-IRF3 protein and IFNβ RNA levels in both cell types (Fig. 4). These data demonstrate that by preventing STING degradation, IFI207 promotes enhanced immune signaling and cytokine responses.

**Fig. 4.**
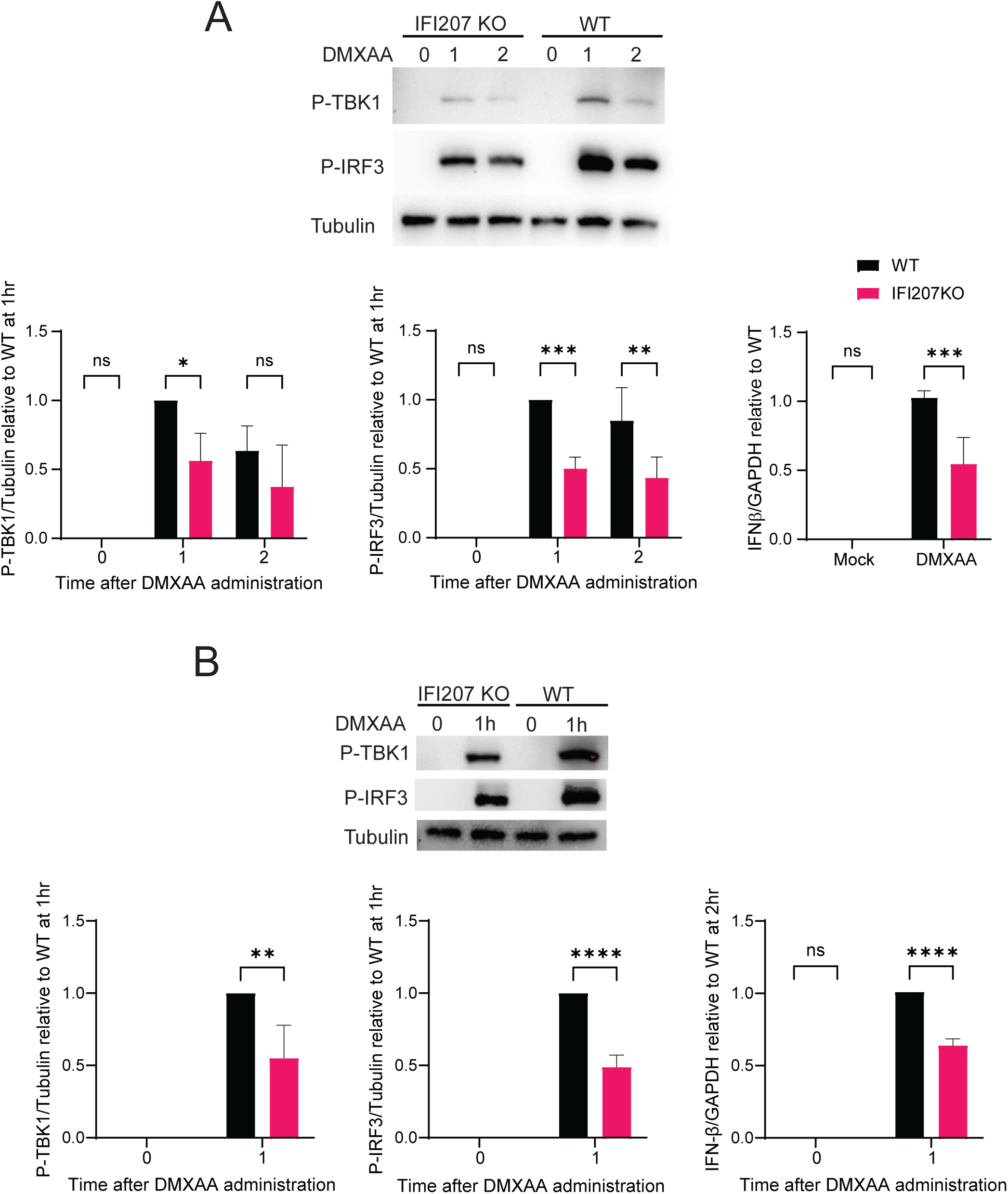
IFI207 enhances P-TBK1, P-IRF3 and type 1 IFN levels. A) BMDMs and BMDCs (B) from the indicated mice were treated with 100 µg/mL DMXAA for 1 or 2 hr. P-TBK1 and P-IRF3 protein levels were quantified by WB. IFNβ mRNA levels following DMXAA stimulation for 2 hr were quantified by RT-qPCR. Shown below each blot is the average of 3 independent experiments ± SD. Two-way ANOVA was used to determine significance. *, *P*≤0.02; **, *P*≤0.004; ***, *P*≤0.0008; ****, *P*≤0.0001 ns, not significant.

There have been many reports regarding the recognition and regulation of DNA viruses by the STING pathway during infection (32, 33). We next investigated whether IFI207 levels affected STING and IFN-1 levels in response to HSV-1 infection; BMDCs from STING mutant (STING^mut^) mice were used as a control. By 6 hpi, the levels of phosphorylated STING (P-STING) and IFN-I were significantly lower in IFI207 KO cells compared to wild-type (WT) cells after incubation (Fig. 5A). Moreover, there was significantly less IFNβ RNA produced in IFI207 KO cells at 6 hr post-infection (Fig.5B). STING^mut^ cells produced almost no IFNβ RNA. Similar results were obtained with BMDMs and fibroblasts in response to HSV-1 infection (Supplementary Fig. 4A). These results indicate that IFI207 promotes DNA virus-triggered innate immune responses.

**Fig. 5.**
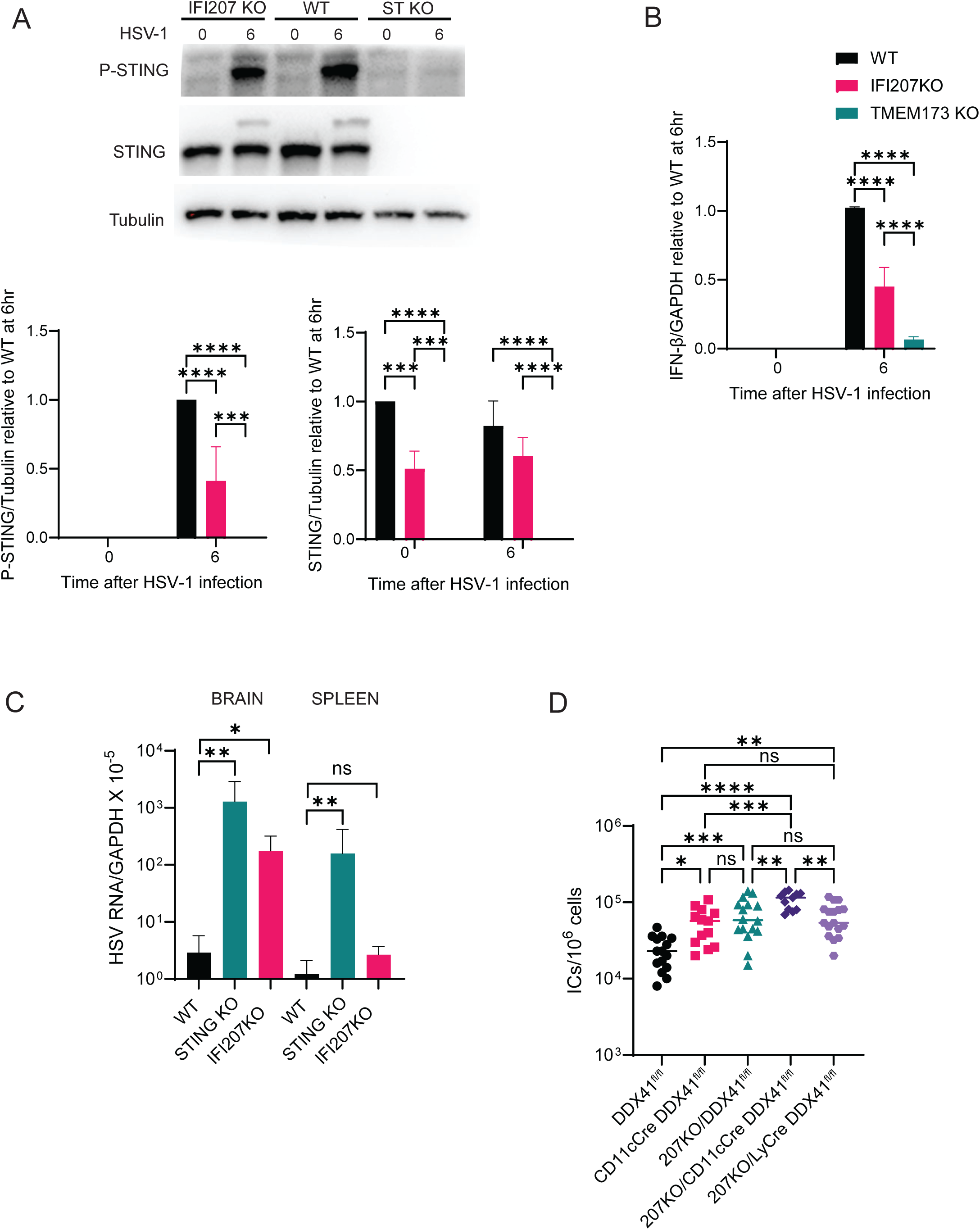
IFI207 promotes STING signaling in response to virus infection. A) and B) BMDMs (A) and BMDCs (B) from the indicated mice were infected HSV-1 for 6 hr (MOI=5), and western blots examining STING and phospho-STING levels were measured. Shown to the right is the quantification of three independent experiments. Two-way ANOVA was used to determine significance. ***, *P*≤0.0008; ****, *P*≤0.0001 B) IFNβ mRNA levels following HSV-1 infection were quantified by RT-qPCR. Shown is the average of 3 experiments. Two-way ANOVA was used to determine significance. ****, *P*≤0.0001. (C) HSV DNA levels in the olfactory bulbs of wild type or IFI207 KO mice 6 days after intranasal inoculation of HSV McKrae. were quantified by qPCR. Welch’s t test was used to determine significance. **, *P*≤0.004. D) Pups of the indicated genotype were infected with 25,000 PFU of MLV. At 16 dpi, splenic viral titers were determined on NIH3T3 cells by infectious center (IC) assays. Each dot represents an individual mouse. Significance was determined by one-way ANOVA. *, *P*≤0.05; **, *P*≤0.001; ***, *P*≤0.00003; ***, *P*≤0.0001

### IFI207 restricts virus infection in vivo

We also infected IFI207 KO, STING mutant and WT mice with HSV-1, to determine if STING stabilization was important for control of virus infection *in vivo*. Six-to-eight-week-old wild type C57BL/6N and IFI207 KO mice received intranasal inoculation of HSV. By 6 dpi, many of the IFI207 KO and wild type mice had lost weight; although not statistically significant, more IFI207 KO mice showed greater weight loss than did WT mice (Supplementary Fig. 4B). We then examined HSV DNA levels in the olfactory bulbs of the mice. The IFI207 mice were more highly infected than the wild type mice, suggesting that STING stabilization by IFI207 *in vivo* leads to better control of HSV infection (Fig. 5C).

We showed previously that MLV infection levels were higher in IFI207 KO mice than in WT mice (23). The DEAD-box helicase DDX41, which recognizes RNA/DNA hybrids generated during MLV reverse transcription, also controls the STING-mediated innate immune response to MLV. Moreover, knockout of DDX41 in the dendritic cell but not macrophage compartment resulted in higher infection levels (34). As we have not generated conditional IFI207 KO mice, to test whether IFI207 exerted its effect in DCs *in vivo*, we crossed both CD11cCre and LysMCre mice with a knocked-in floxed DDX41 allele with IFI207 KO mice to generate DC and macrophage-specific IFI207 KO/DDX41^fl/fl^ double KO mice, respectively. Newborn mice were injected with MLV, and splenic viral titers were measured at 16 days post-infection. DDX41^fl/fl^ mice (without Cre), IFI207 KO/DDX41^fl/fl^ and CD11c /DDX41^fl/fl^ mice were included. As we observed previously, deletion of IFI207 alone (207-/- DDX41fl/fl) or DDX41 alone in DCs (CD11cCre+ DDX41fl/fl) resulted in higher infection levels than in WT (DDX41fl/fl) mice (Fig. 5D) (23, 34). Knockout of DDX41 in DCs in combination with IFI207 knockout (207-/- CD11cCre+ DDX41fl/fl) but not in macrophages (207-/-LyCre DDX41fl/fl) resulted in the highest infection levels (Fig. 5D). This suggests that IFI207 stabilization of STING in DCs, which are the initial targets of MLV infection *in vivo*, is important for the control of infection.

## Discussion

The innate immune system serves as a first line of defense by utilizing a variety of PRRs that detect various nucleic acid PAMPs generated during virus infection, many of which activate STING and its downstream pathways (35). Among the intracellular DNA sensors that act upstream of STING, cGAS is the most extensively studied (36). Another sensor, DDX41, recognizes both DNA and DNA/RNA hybrids generated upon retrovirus infection and virus-induced mitochondrial damage and activates the STING pathway (34, 37, 38). Many studies have also implicated various ALRs. Notably, IFI16 in humans responds to DNA from HSV and other viruses during infection (39–41). Several mouse ALRs, including IFI203 and IFI204 are also thought to sense viral DNA (42, 43).

In contrast, we show here that rather than functioning as a PRR, IFI207 enhances innate immune responses by stabilizing STING protein after activation by cGAMP, the agonist DMXAA, synthetic dsDNA, the retrovirus MLV and HSV-1, leading to an increased antiviral response [here and (23)]. While our previous study showed that IFI207 protected STING from degradation, the molecular mechanisms by which this occurred were unknown. In the present study, we show that IFI207 promotes antiviral responses by modulating K63 linked ubiquitination of STING and subsequent binding to HRS, thereby inhibiting its degradation via the ESCRT machinery.

Post-translational modifications such as ubiquitination, phosphorylation, palmitoylation, SUMOylation, acetylation and glutamylation are all involved in STING regulation (44, 45). STING activation can be positively or negatively regulated by K6-, K11-, K27-, K48-, and K63-linked polyubiquitination (19, 44–46). In particular, TRIM32 and TRIM56 target STING for K63-linked ubiquitination and this plays a role in the defense against virus infections (16, 17).

The ESCRT machinery also plays a role in STING degradation following its activation (10, 11, 21). HRS, a key component of the ESCRT-0 complex, specifically recognizes K63-linked ubiquitinated proteins, such as STING, and recruits them to acidic endosomes (10, 21). Ubiquitinated STING forms a platform in the endosomal compartment for the recruitment of the ESCRT complex, and its association with the complex promotes STING degradation through fusion of vesicles coated with oligomeric STING with the endolysosome (21). Here we show that the ALR IFI207, which is under strong positive selection, modulates the antiviral innate immune response via binding to and protecting activated STING from K63 ubiquitination, and subsequent interaction with HRS and degradation via the ESCRT machinery. Thus, IFI207/STING interaction may regulate the turnover of STING by ensuring that sufficient levels are available for rapid signal transduction upon pathogen infection. IFI207 also appears to be degraded in cells activated by DMXAA, although this degradation is not rescued by TAK243, suggesting that this is ubiqutination-independent (Fig. 2A and 3C). One possibility is the IFI207 in complex with STING and HRS is also transported to the degradative compartment. This may represent a negative feedback loop that would limit the enhancing effect of IFI207 after STING activation.

We do not yet know which of the critical K63 residues is protected from ubiqutination by IFI207; our preliminary data suggest that more than one residue may play a role. Although IFI207 is unique to mice, it is likely that other host proteins in other species carry out this function. IFI16, although it has a single copy of the S/T-rich repeat region found in IFI207, binds STING via its pyrin domain, which is thought to lead to IFI16 degradation (23, 47) (Fig. S2). Thus, IFI207’s putative coiled domain may be the means by which it protects STING from degradation.

We previously showed that DDX41 was required to activate the full STING-dependent anti-viral response to MLV and that DDX41 expression specifically in DCs contributed to effective *in vivo* control of MLV infection (34). To test whether the DDX41-dependent STING response also was altered by loss of IFI207 in vivo, we generated mice with a DC-specific floxed allele of DDX41 on an IFI207 knockout background. Mice that lacked both DDX41 and IFI207 in DCs were more susceptible to MLV infection than were mice lacking DDX41 in DCs or IFI207 alone, suggesting that IFI207-mediated stabilization of STING in DCs, the initial targets of MLV infection *in vi*vo, is critical in the control of retrovirus infection.

It may be that murine retroviruses, which have existed in mice for millions of years, or other viruses such as herpesviruses, were responsible for the selective pressure that caused the high level of diversification of the ALR locus in *Mus*. Here we show that IFI207 promotes HSV-1-triggered innate immune responses in DCs, fibroblasts and macrophages and protects mice from infection. While Mus species do not harbor any known alphaviruses like HSV-1, they do harbor the betaherpesvirus Murine Cytomegalovirus and they can be infected with Murid gammaherpesvirus 68 (MHV-68) (48). Interestingly, we found previously that the only other species besides *Mus* to harbor an *IF207*-like gene was *Apodemus*, although it only encodes two S/T- rich repeats (23). *Apodemus* rodents are known to be infected with MHV-68 and similar gammaherpesviruses, so it is tempting to speculate that herpesviruses could play a role in IFI207’s positive selection (49).

In conclusion, our findings indicate a novel mechanism by which STING-mediated antiviral innate immune responses triggered by DNA viruses and MLV may be regulated. Following infection, IFI207 interacts with STING, inhibiting K63-linked ubiquitination, thereby preventing STING from binding to HRS and degradation through the ESCRT pathway (Fig. 6). Thus, the impact of IFI207 on STING ubiquitination is crucial for the regulation of STING translocation and subsequent STING-mediated antiviral responses. As the STING pathway is downstream of several nucleic acid sensors, including cGAS, ALRs, and DDXs, elucidating the precise mechanisms that regulate STING signaling through the STING degradation is becoming increasingly important for understanding various diseases in which controlling infection and suppressing aberrant STING-dependent inflammation can be beneficial. The outcomes of this research are expected to contribute to the development of innovative therapeutic and preventive strategies against viral infections.

**Fig. 6.**
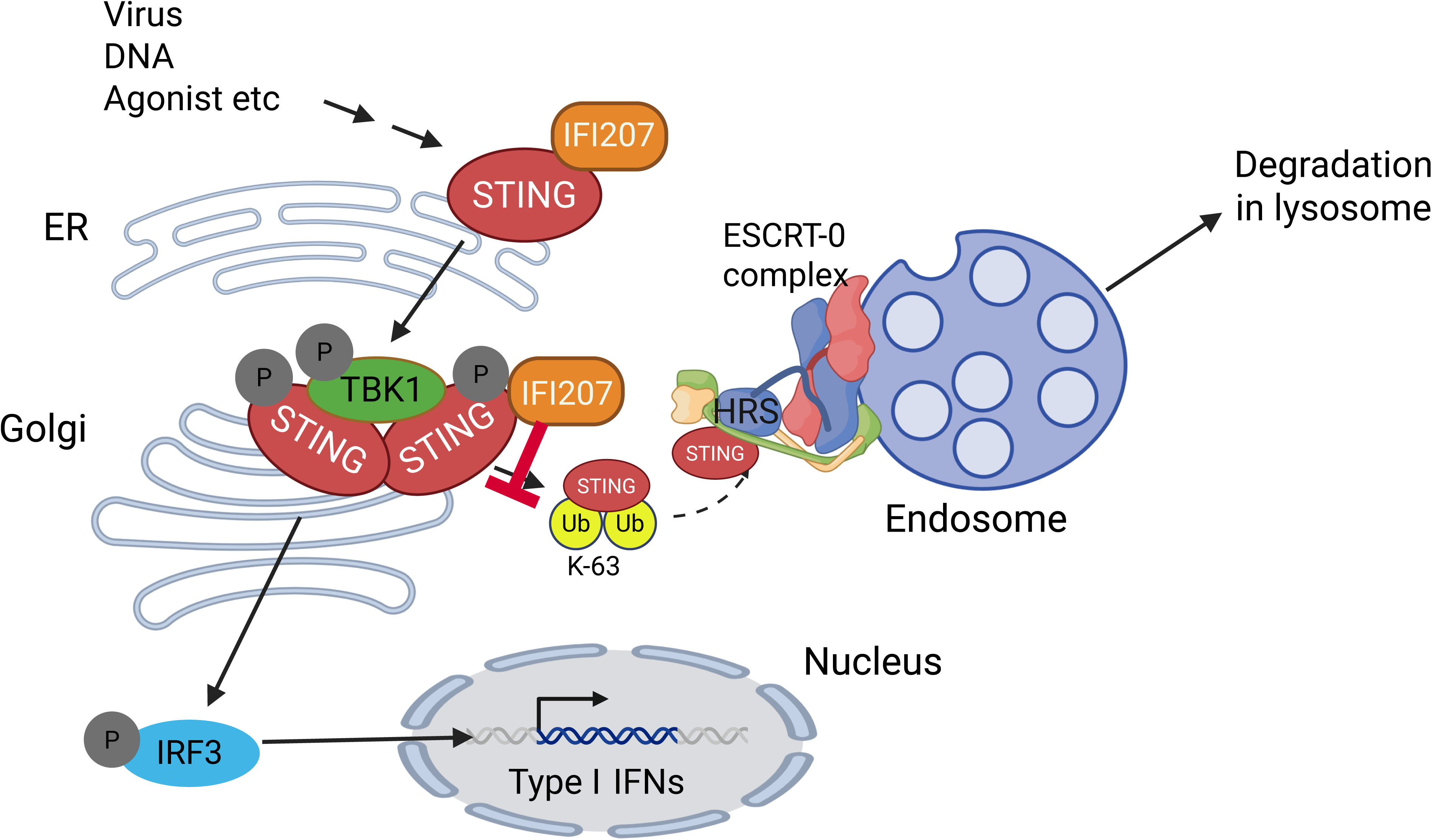
Model of IFI207’s inhibition of STING degradation. STING is bound to IFI207, and after activation by DNA, dimerizes and moves from the ER to the Golgi. After the recruitment of TBK1 and STING phosphorylation, it normally becomes K63-ubiquitinated, binds HRS and is escorted to the endosome and then to the lysosome, where it is degraded. When bound to IFI207, K63 ubiquitination is inefficient, STING levels are higher, and STING downstream activation is increased, leading to a stronger type I IFN response. *Created in BioRender. ENYA, T.* (*2025*) https://BioRender.com/3yt291e.

## Materials and Methods

### Ethics Statement

All mice were housed according to the policies of the Animal Care Committee (ACC) of the UIC; all studies were performed in accordance with the recommendations in the Guide for the Care and Use of Laboratory Animals of the National Institutes of Health. The experiments performed with mice in this study were approved by the UIC ACC (protocol no. 21-125).

### Mice

IFI207 KO, DDX41^fl/fl^, CD11cCre DDX41^fl/fl^ and LysMCre DDX41^fl/fl^ mice were previously described (23, 34). 207KO/DDX41^fl/fl^, 207KO/CD11cCre DDX41^fl/fl^ and 207KO/LycCre DDX41^fl/fl^ mice were generated by backcrossing. C57BL/6N, LysMCre (B6.129P2-Lyz2tm1(cre)Ifo/J), CD11cCre (B6.Cg-Tg(Itgax-cre)1-1Reiz/J) and STING^mut^ (C57BL/6J-Sting1gt/J) mice were originally purchased from The Jackson Laboratory and were bred at the University of Illinois at Chicago (UIC). In some experiments, DDX41^fl/fl^, CD11cCre and LysMCre mice were used as WT controls.

### Cell culture

Bone marrow was harvested from hind limbs of mice. Primary BMDMs and BMDCs were cultured, collected and analyzed by flow cytometry as previously described (23). BMDMs and BMDCs cell populations were >80% pure, as determined by flow cytometry and staining with anti- CD11b, or -CD11c antibodies, respectively. Fibroblasts were isolated from mouse tails and cultured in Dulbecco’s Modified Eagle Medium (DMEM; Invitrogen) containing 10% fetal bovine serum (FBS; Invitrogen), 2 mM L-glutamine, 100 U/ml penicillin, 100 ug/ml streptomycin in 5% CO2 at 37°C. Fibroblasts were used for experiments at passage 2 or 3. HEK293T and NIH3T3 cells were cultivated in the same media at 37°C in 5% CO2 at 37°C. Cells were treated with DMXAA at 100μg/ml (Sigma Aldrich, D5817) and harvested at the indicated times for protein and RNA analyses.

### Plasmids and transfections

The V5-tagged IFI207 and FLAG-tagged STING plasmids were previously described (23, 25). The HA-tagged Ub-WT HA-Ub (#17608) and HA-Ub (K63 only) (#17606) plasmids were from Addgene. HEK293T cells were transfected with expression plasmids using Lipofectamine 2000. When different amounts of plasmids were transfected within experiments, empty vector (EV) plasmid [pcDNA 3.1/Myc (+)] was added to keep the total amount of transfected DNA equal for all samples. Twenty-four hrs after transfection, the cells were collected for protein expression analysis.

### Immunoprecipitations

HEK293T cells were transfected with expression plasmids for 24 hr using Lipofectamine 2000, as described in the section Transfections for Protein Expression Analysis. The cells were lysed in radioimmunoprecipitation (RIPA) buffer (50mM Tris pH 8, 150mM NaCl, 1mM EDTA, 1% Triton X-100, 1% sodium deoxycholate, and 0.1% SDS) containing protease and phosphatase inhibitor cocktail (Thermo Scientific 78443), sonicated, and quantified by Bradford Assay (Bio-Rad). For ubiquitination assays, the cells were lysed in RIPA buffer, supplemented with the protease and phosphatase inhibitor cocktail and 20 μM of DUB inhibitors PR-619 (LifeSensors SI-9619). For immunoprecipitation (IP) assays, the extracts were incubated with the indicated primary antibodies for 2-4 hrs, and Protein A/G PLUS-Agarose (Santa Cruz sc-2003) was added and incubated at 4°C overnight on a rotating rack. IP samples were washed four times in RIPA buffer and were centrifuged for 3 min (2,500 rpm) and eluted in 4X Laemmli buffer containing 5% β-mercaptoethanol. For IP assays with primary BMDMs, the cells were lysed in NP40 lysis buffer (40 mM Tris–HCl [pH 7.4], 2 mM EDTA, 2% Nonidet P40 [IGEPAL], 20% Glycerol) containing the protease and phosphatase inhibitor cocktail.

### Western blots

For Western blot analyses, proteins were extracted in RIPA buffer containing fresh protease and phosphatase inhibitor cocktail (Thermo Scientific 78443). The lysates were sonicated on ice, quantified by Bradford Assay, prepared in 4X Laemmli buffer plus 10% β-mercaptoethanol, and boiled for 5 mins. Proteins from cell lysates were resolved by sodium dodecyl sulfate polyacrylamide gel electrophoresis (SDS-PAGE) and then transferred to polyvinylidene difluoride membranes (Millipore IPVH00010). For ubiquitination assays, the cells were lysed in RIPA buffer, supplemented with the protease and phosphatase inhibitor cocktail and 20 μM of DUB inhibitors PR-619 (LifeSensors SI-9619). Proteins were detected with primary antibodies against STING (Cell Signaling Technology [CST] D1V5L), phospho-STING (CST D8F4W), phospho-TBK1 (CST D52C2), phospho-IRF3 (CST 4D4G), HA-tag (CST C29F4), FLAG-tag (Sigma F1804), V5-tag (Invitrogen 46-0705) or α-Tubulin (Sigma T6199). HRP-conjugated antibodies (CST anti-rabbit or anti-mouse HRP IgG,7074 and 7076, respectively) were used for detection with Pierce ECL Western Blotting Substrate (Thermo Fisher Scientific, 322209), ECL Prime Western Blotting Reagent (GE Healthcare Amersham, RPN2232) or SuperSignal West Femto Maximum Sensitivity Substrate (Thermo Fisher Scientific, 34095).

### Quantitative RT-PCR

RNA was isolated by RNeasy Mini Kit (QIAGEN 74106) with on-column DNase digestion using RNase-Free DNase Set (QIAGEN 79254). cDNA was synthesized with random hexamers from the Superscript III First-Strand Synthesis System (Invitrogen 18080051). RT-qPCR was performed using Power SYBR Green PCR Master Mix (Applied Biosystems 4367659) with the indicated primer sets on a QuantStudio 5 Real-Time PCR System (Applied Biosystems) (Table 1).

**Table 1.**
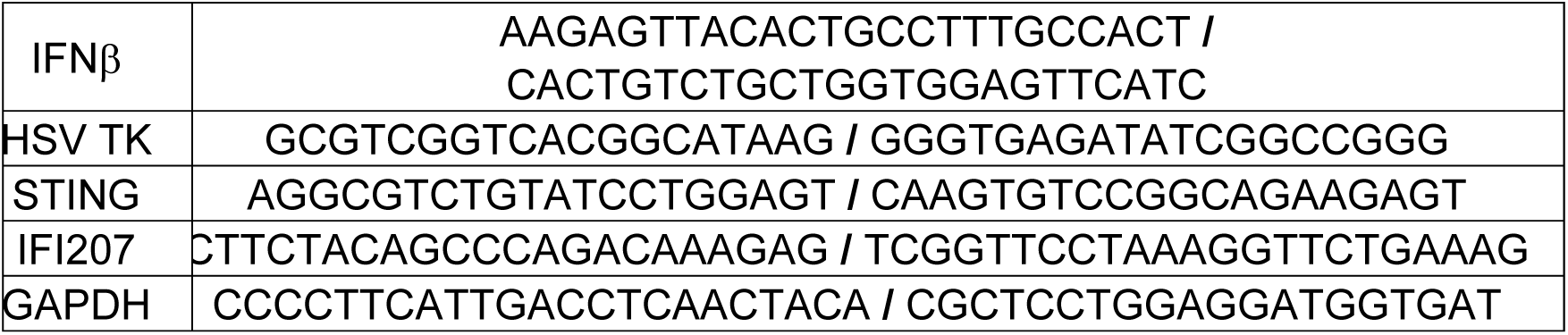
Primers used for RTqPCR and DNA qPCR.

### ELISAs

For analysis of interferon secretion, fibroblasts were seeded into 12-well plates. The next day, cells were pre-treated with BafA1 or TAK243 for 30 minutes, followed by stimulation with DMXAA for 3 hours. Cell culture supernatants were then collected, and IFN-β levels were measured using the LumiKine Xpress mIFN-β 2.0 (InvivoGen, Cat# luex-mifnbv3) according to the manufacturer’s instructions.

### In vitro HSV-1 infection of primary BMDMs and BMDCs

HSV-1 (KOS strain; ATCC VR1493) was used for in vitro experiments. BMDMs and BMDCs from WT, IFI207KO and STING^mut^ mice were infected with HSV-1 at multiplicity of infection (MOI) of 5 in media containing 2% FBS. Two hrs after infection, the virus was removed, and fresh medium was added. Six hrs after infection, the cells were washed with PBS and collected for RNA isolation and western blot analysis.

### MLV purification and titer

Moloney MLV were isolated from supernatants of stably infected NIH3T3 cells, and virus titers were determined by an infectious center (IC) assay on NIH3T3 cells using monoclonal anti-ENV (Ab538) antibody, as previously described (50). ICs were counted by microscopy (Keyence BZ-X710 and BZ-X analyzer).

### In vivo infections

Two-day old mice of the indicated genotype were injected intraperitoneally with 2.5 × 10^4^ PFU MLV. At 16 days post-infection, splenocytes were harvested and treated with ammonium chloride-potassium buffer. Tenfold dilutions of the cells were co-cultured with NIH3T3 cells for 3 days to measure viral titers by IC assays.

For HSV, six- to eight-week-old mice IFI207 KO and wild type mice received 10^7^ PFU of HSV McKrae via intranasal inoculation. The mice were monitored daily and at 6 days post-infection, the olfactory bulbs were collected and used to extract DNA. The primers were used for RTqPCR to measure infection levels are listed in Table 1. All reactions were normalized to GAPDH.

### Statistical analysis and data availability

Data shown are the averages of at least 3 independent experiments, or as otherwise indicated in the figure legends. Unpaired one- and two-tailed t tests were performed using GraphPad Prism 10.1 software to calculate *P* values. All raw data have been deposited in the Mendeley data set found at https://data.mendeley.com/preview/4r6sffxsz2?a=f07ae714-2922-4afa-b5d0-22ff4a87a6d5.

## Acknowledgments

We thank David Ryan for help with the mouse breeding and setting up the genetic crosses, and the members of our lab for helpful discussions. Molecular graphics and analyses were performed with UCSF Chimera, developed by the Resource for Biocomputing, Visualization, and Informatics at the University of California, San Francisco, with support from NIH P41-GM103311 (51). This study was supported by the National Institutes of Health Grants R01AI121275 and 1R01AI174538 (S.R.R.).

---

**Suppl. Fig. 1.**
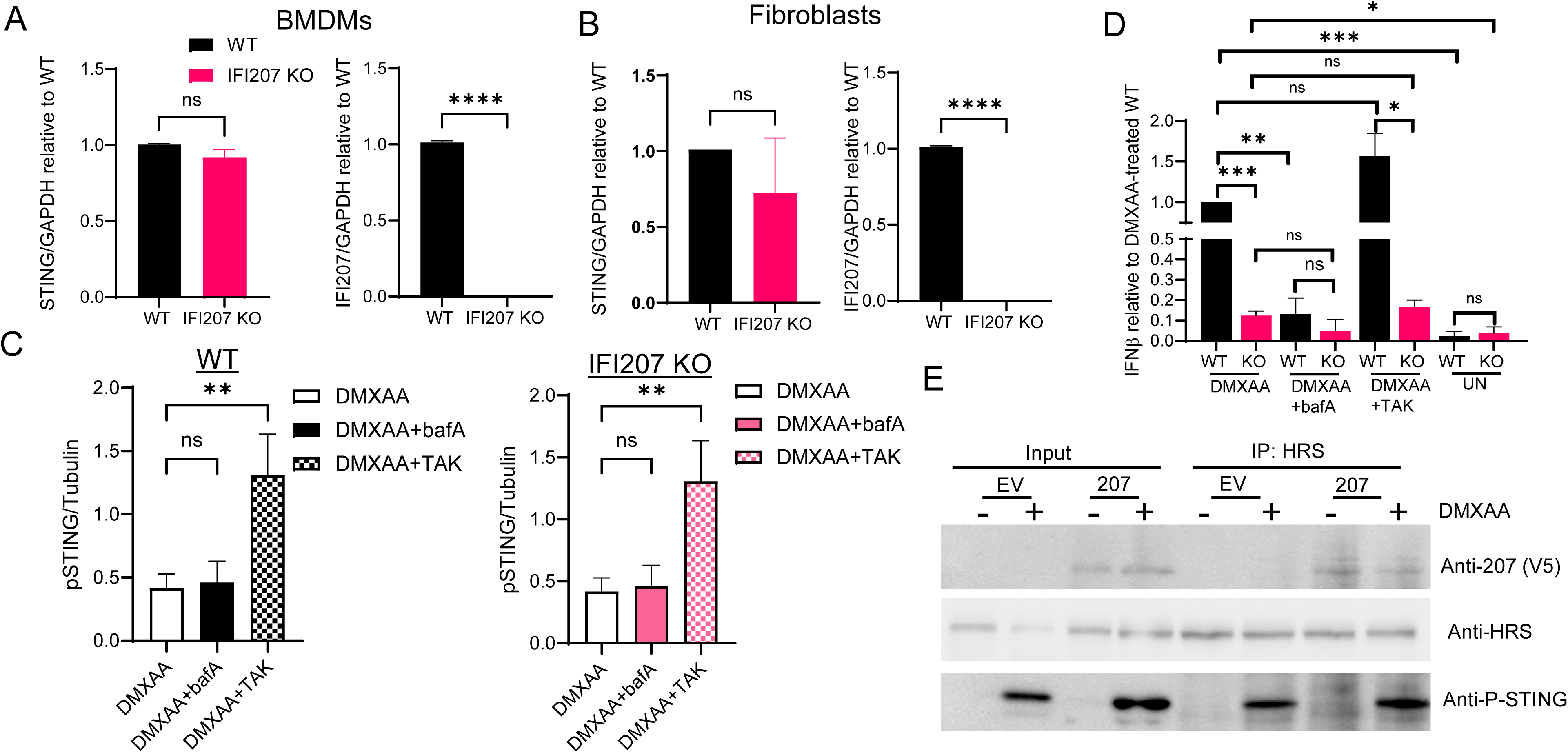
IFI207 promotes STING signaling. BMDMs (A) and fibroblasts (B) from IFI207 KO and wild-type mice were treated with 100 µg/mL DMXAA and RNA was examined by RT-qPCR for the levels of STING and GAPDH. Quantification of 3 independent experiments ± SD is shown. Paired t-tests were used to determine significance. ****, *P*≤0.007. (C) Comparison of p-STING levels in DMXAA, DMXAA+bafA and DMXAA-treated fibroblasts from WT and IFI207 KO mice 3 hr post-DMXAA treatment. Replotted data from Fig. 1A. One-way ANOVA was used to determine significance. **, *P*≤0.005. (D) IFNα levels produced in WT and IFI207 KO fibroblasts was measured by ELISA at 2 hr post-DMXAA treatment. UN, untreated. Quantification of 3 independent experiments ± SD is shown. Two-tailed t-tests were used to determine significance. *, P≤0.05, **, P≤0.01, ***, P≤0.001.(E) HRS co-immunoprecipitates with pSTING.

**Suppl. Fig. 2.**
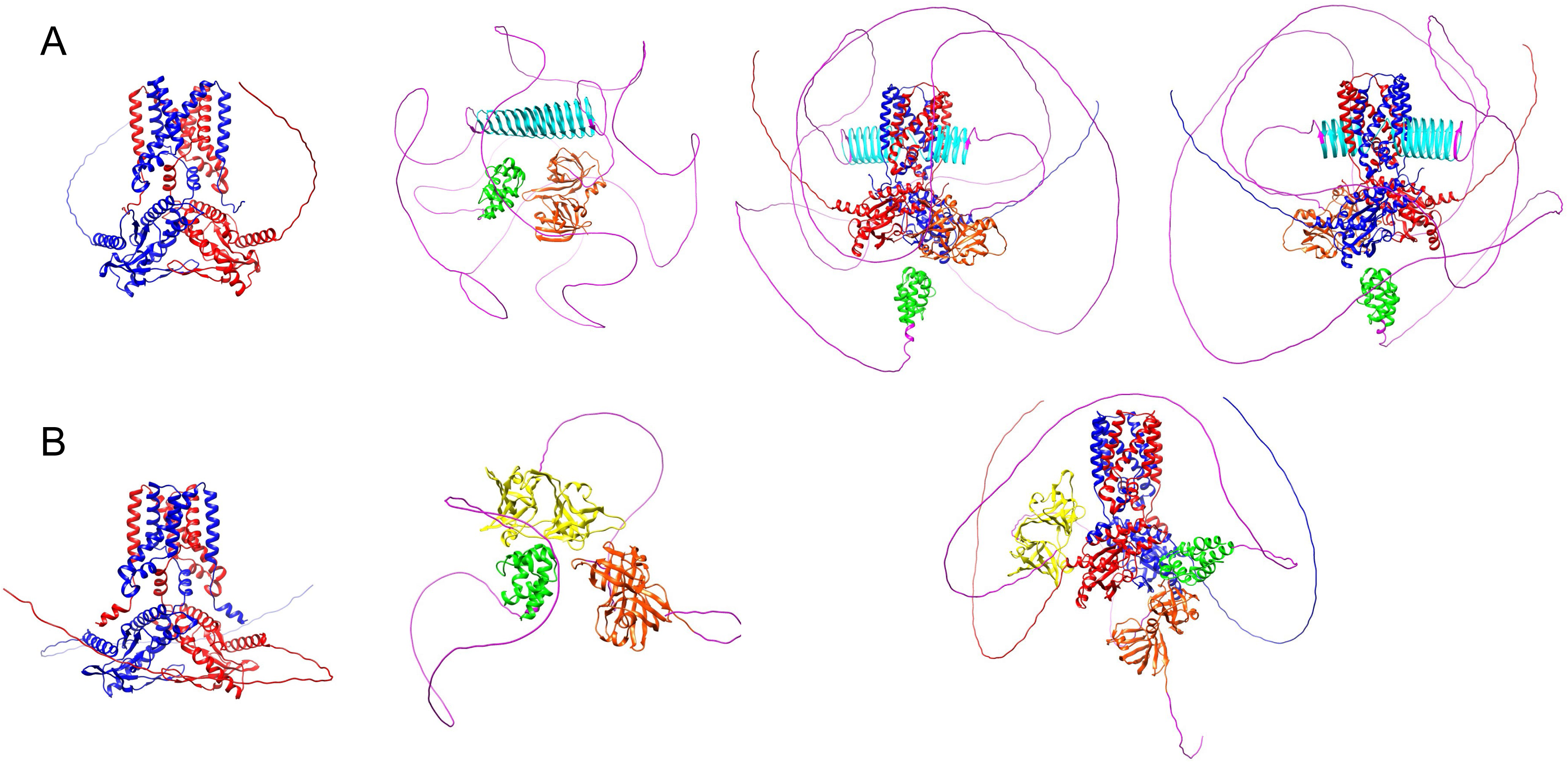
A) AlphaFold predicted structure of the mouse STING dimer (red and blue) and IFI207 (magenta). The mouse STING/IFI207 complex is shown in both front and back. The N terminal pyrin domain is green, the C terminal HINB domain is orange, the repeat region is cyan. B) AlphaFold predicted structure of the human STING dimer (red and blue) and IFI16 (magenta). Pyrin and HINB domains colored as in A; the HINA domain is yellow. Structures were downloaded from AlphaFold Server (https://alphafoldserver.com/) (Abramson, 2024). UCSF Chimera was used to generate images (https://www.rbvi.ucsf.edu/chimera) (Petterson et al., 2004).

**Suppl. Fig. 3.**
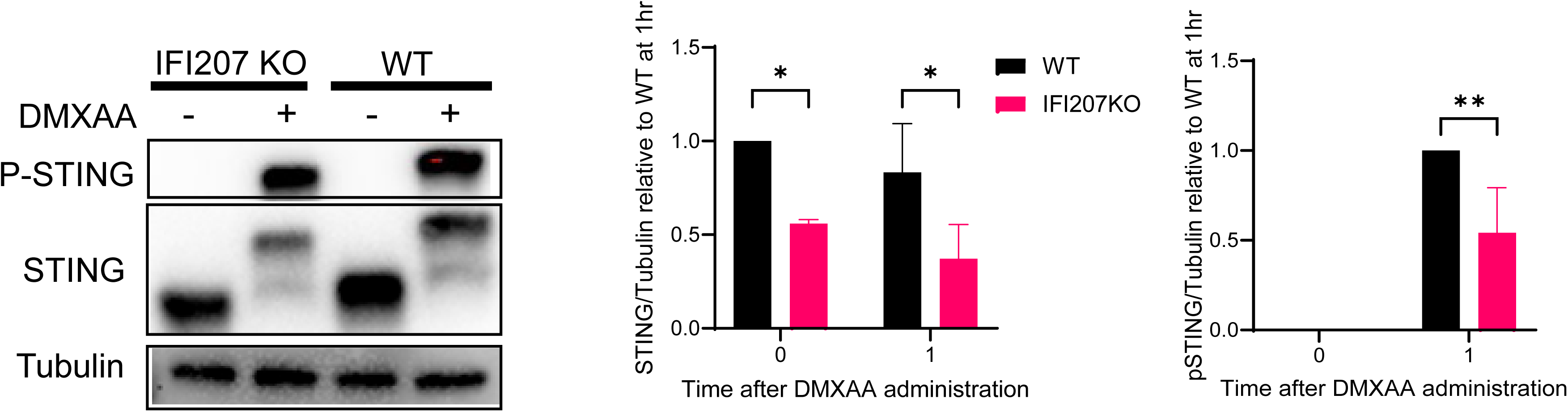
IFI207 stabilizes STING in BMDCs. BMDCs from IFI207 KO and wild-type mice were treated with 100 µg/ml DMXAA and STING and pSTING protein levels were examined by western blot at 1 hr post-treatment. Shown to the right is quantification of 3 independent experiments ± SD. Two-way ANOVA was used to determine significance. *, *P*≤0.02; **, *P*≤0.004.

**Suppl. Fig. 4.**
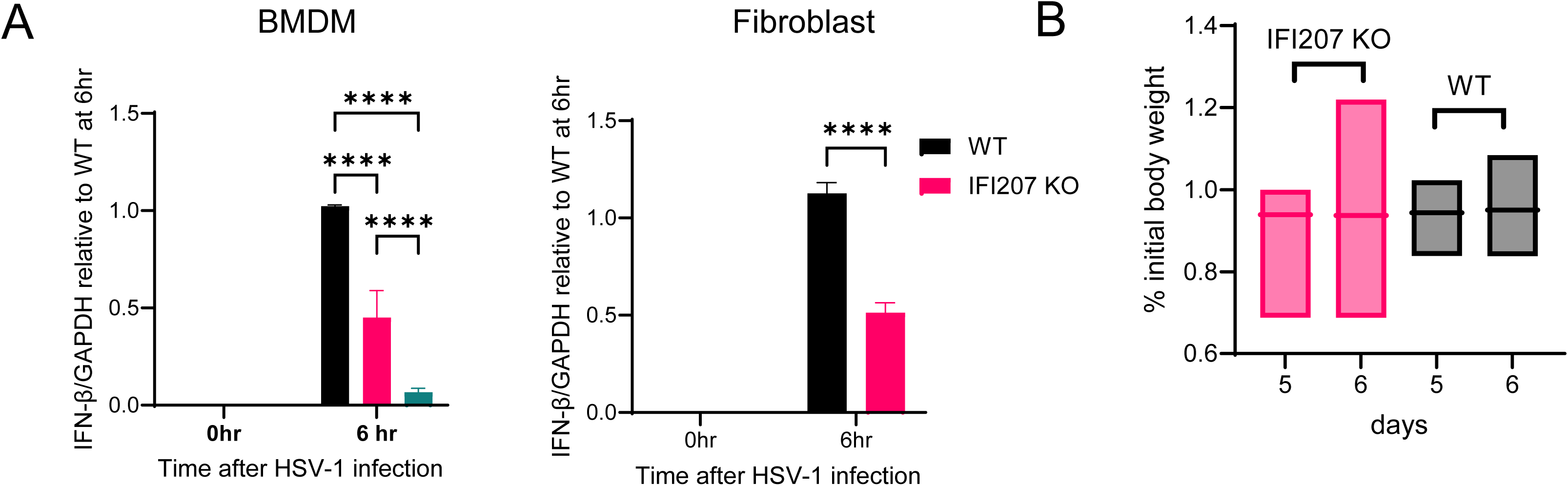
IFI207 promotes STING signaling in response to virus infection. A) BMDMs and fibroblasts from the indicated mice were infected HSV-1 for 6 hr (MOI=5), IFNβ mRNA levels were quantified by RT-qPCR. Shown is the average of 3 experiments. Two-way ANOVA was used to determine significance. ****, P≤0.0001. B) average weight loss in IFI207 KO and wild type mice at 5- and 6-days post-infection.

